# Analysis of sugar-induced cell death dynamics in S. cerevisiae strains with deleted genes involved in several key metabolic processes

**DOI:** 10.1101/2023.11.10.566565

**Authors:** Airat Ya. Valiakhmetov

## Abstract

**Background:** incubation of exponentially growing yeast S.cerevisiae with glucose in the absence of other nutrients results in **S**ugar **I**nduced **C**ell **D**eath (SICD). SICD is accompanied by the accumulation of reactive oxygen species (ROS), has the nature of primary necrosis, affects cells in the S-phase of the cell cycle, and is completely suppressed by dissipation of ΔΨ. The specific mechanism linking the ΔΨ status to the induction of SICD remains unclear. This study aimed to attempt to identify the specific molecular mechanism responsible for ROS overproduction and the development of SICD.

**Methods:** The main method employed was the analysis of SICD development in a set of knockout mutants targeting key participants in metabolic processes.

**Results:** A statistically significant decrease in the number of cells with ROS overproduction was observed in the ΔAFO1, ΔPOX1, ΔYNO1, ΔTRK1, ΔTRK2, ΔVSB1, and ΔYPR003C strains. A significant decrease in the number of cells with SICD was shown in the ΔTRK1, ΔVSB1, and ΔYPR003C strains. The development of SICD is not due to the presence of a nitrogen reactive species (NRS). Deletion of certain genes expressed during the S-phase of the cell cycle did not alter the dynamics of ROS accumulation and the development of SICD. The presence of exogenous or endogenous glutathione significantly suppresses both processes studied, although not as effectively as ΔΨ dissipation.

**Conclusions:** The development of SICD is dependent on the presence of ROS, but is not strictly linked to it, as evidenced by the effects of glutathione and mutations related to its biosynthesis. In all strains tested, SICD was critically dependent on ΔΨ, although the nature of its generator remains unclear.

## 1 Introduction

The life cycle of an individual cell ends with death. After the discovery of apoptosis, cell death was no longer considered a spontaneous and unregulated event. It is currently believed that apoptosis and autophagy are types of cell death that are regulated and programmed. This type of death is characterized by a high degree of temporal organization in the processes of cellular structure decay, which culminate in the rupture of the cell membrane - secondary necrosis. In contrast, necrosis is considered an unregulated type of cell death, triggered by extreme external factors and starting with the rupture of the plasma membrane - primary necrosis [1]. Apoptosis was long considered characteristic only of metazoans until 1997 when a study describing apoptotic markers in S. cerevisiae yeast was published [2]. This discovery served as the impetus for a more careful analysis of the phenomenon of death in unicellular organisms. Apoptosis in yeast has been described in many cases [3-7]. Of particular interest to us is SICD - the death of stationary yeast cells incubated with glucose but in the absence of other essential nutrients for growth. Initially described in 1991 [8], SICD was later found to have the nature of apoptosis and was induced by an excess of ROS [9]. Further studies showed that SICD is independent of ACP pathways [8] and involves glucose or fructose phosphorylation steps [10,11]. Interestingly, SICD has also been observed under strictly anaerobic conditions in the bottom-fermenting yeast Saccharomyces pastorianus [12]. In an attempt to elucidate the specific mechanism underlying SICD, attention was drawn to the Crabtree effect [13, 14]. It is hypothesized that the Crabtree effect involves competition between the mitochondrial respiratory chain and glycolytic enzymes for ADP and inorganic phosphate. The suppression of SICD in stationary yeast cultures by exogenous phosphate and succinate, according to the authors, confirms this assumption. It is important to note that all these studies were conducted with yeast in the stationary growth phase. Later, we showed that incubation of exponentially growing yeast S.cerevisiae with glucose in the absence of other components necessary for growth also leads to SICD, which was of the nature of primary necrosis [15]. Unlike stationary cells, SICD in exponentially growing yeast occurred much faster (minutes vs. days for cells in the stationary phase) as a result of the nuclear, mitochondrial, and plasma membrane raptures and affected only cells in the S phase of the cell cycle. A notable feature of SICD in growing culture was its sensitivity to extracellular pH and membrane potential. Incubation of yeast with glucose in a medium with pH 7.0 (ΔpH dissipation) or in the presence of 150 mM KCl (ΔΨ dissipation) completely prevented SICD [16]. These facts cast doubt on the thesis that primary necrosis occurs only as a result of extreme external physical or chemical impact on the cell. SICD occurred under conditions where there were no extreme external impacts on the cell. Instead, an imbalance in the availability of all nutrients needed for growth led to SICD in the form of primary necrosis. Notably, the mitochondrial respiratory chain in exponentially growing yeast is not involved in the development of SICD. This was shown both in the case of petite mutants obtained by treating yeast cells with ethidium bromide and in the ΔAFO1 strain, which lacks a large mitochondrial ribosomal subunit [16]. Another feature of SICD in growing yeast is the release of octanoic acid, a highly toxic intermediate of alcoholic fermentation, into the medium [17]. It may be octanoic acid that causes the rupture of both plasma and organelle membranes. However, direct evidence has not yet been received. It is also important to note that no relationship has been identified between ΔΨ status, ROS generation, on the one hand, and metabolic pathways for the synthesis of octanoic acid, on the other. The role of ROS is critical in the development of SICD in growing yeast since we were able to demonstrate the fact of increased ROS content specifically in dead cells [15]. This study focuses on the analysis of some critical elements of cellular machinery that could establish a direct link between ROS generation and SICD. The second task was to identify the node of cellular metabolism that tightly links the development of SICD with the status of the membrane potential on the yeast plasma membrane. The third goal was to screen some genes expressed in the S phase of the cell cycle to identify a possible trigger for SICD.

## 2 Materials and Methods

### 2.1 Culture growth

Strains used in this study are listed in Table 1 and were obtained from the Euroscarf collection. The culture grew on the standard YPD medium (Applichem, Darmstadt, Germany) for 15–17 h (mid-exponential phase) and was twice washed with distilled water. Yeast cells were pelleted and suspended (1 g/10 mL w/w, 1.45 × 10^8^ cells/mL) in MilliQ water.

**Table 1.**
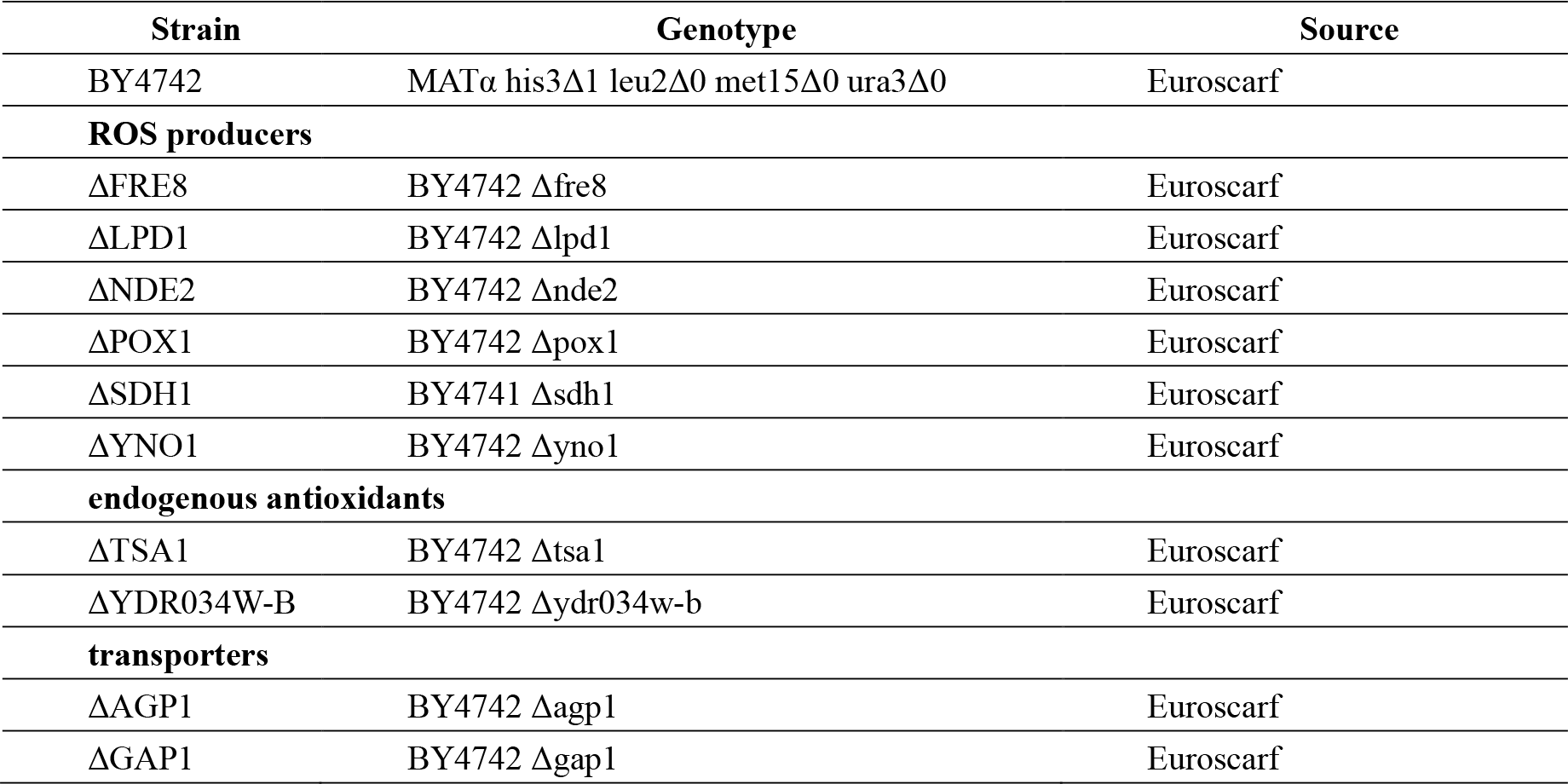
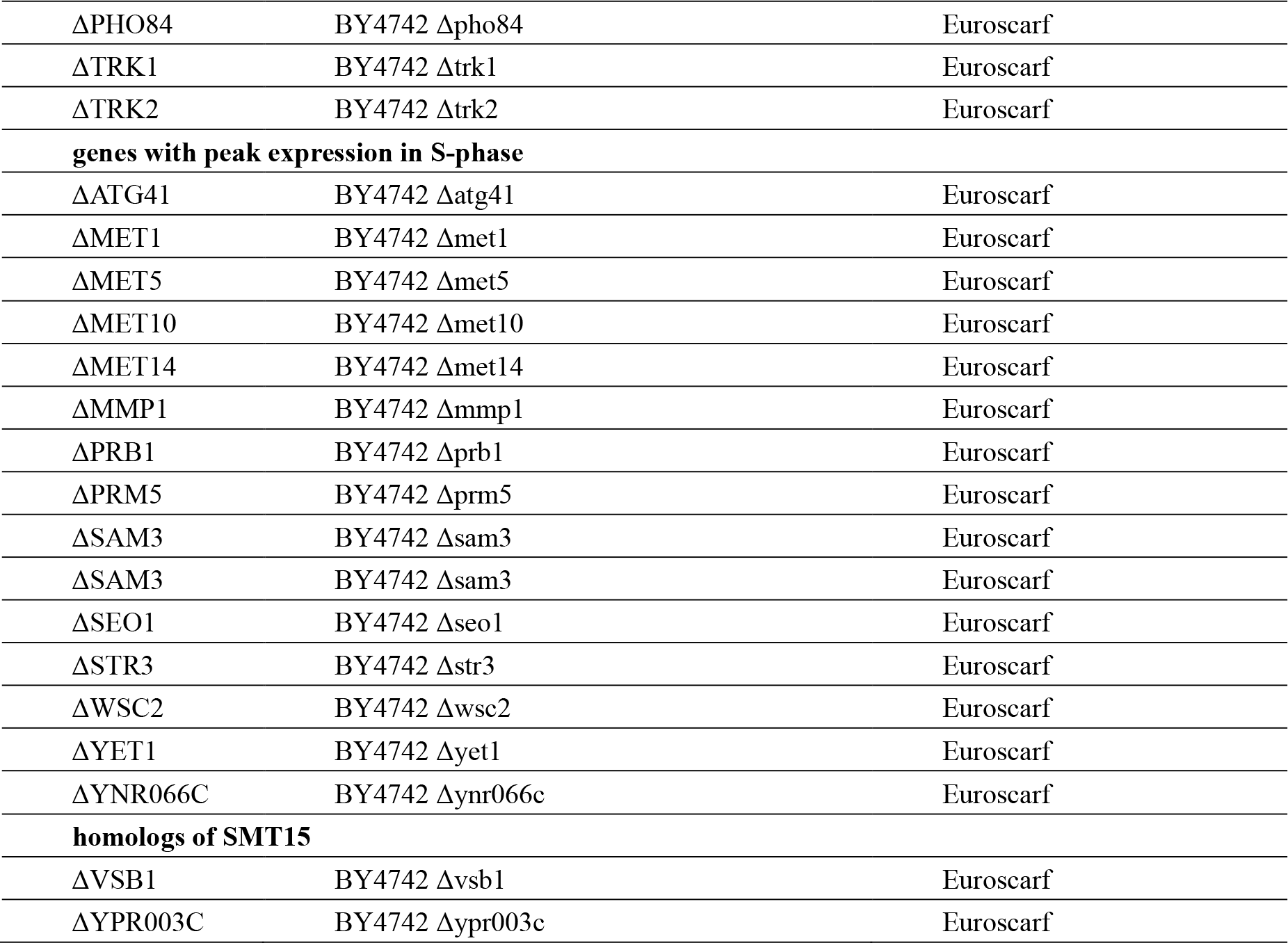
Strains used in this study.

### 2.2 1,2,3,- Dihydrorhodamine (DHR) Staining

To estimate the number of cells with elevated intracellular ROS levels, we used DHR (Sigma-Aldrich, St. Louis, MO, USA). 5 µL of 0.5 mg/mL DHR in DMSO was added to 250 µL cell suspension in water. The samples were incubated in a ThermoMixer (Eppendorf, Hamburg, Germany) for 1 h at 30°. The cells were pelleted at 13,000× g for 1 min, and the supernatant was discarded. The cells were washed once with 0.5 mL of water, and the cell pellet was suspended in 250 µL water. Another 250 µL of cell suspension in water without the addition of DHR was subjected to all the above steps to obtain cells for propidium iodide (PI) staining (see below).

### 2.3 SICD Assay and Flow Cytometry

SICD assay for DHR-stained and unstained cells was done in a parallel series of tubes. 250 µL of cells obtained as described above was distributed in 50 µL into four tubes. This was the standard test kit for the SICD. Nothing was added to the first tube—this was the control. The second tube served as the main sample for checking the SICD. In the third tube, water was replaced with buffer (50 mM HEPES, pH 7.0) to test the effects of neutral pH on the SICD. 2.5 µL of 3 M KCl was added to the 4th tube to check the value of the membrane potential on SICD. Then, to induce SICD, 2.5 µL of 2 M glucose solution was added to all tubes (final [C] = 100 mM), except for the 1st one. The samples were incubated in a ThermoMixer (Eppendorf, Hamburg, Germany) at 30 °C for 1 h. To proceed to flow cytometry to the samples with DHR-stained cells 1 mL of water was added. PI staining was used to determine the percentage of dead cells. To the samples with not DHR-stained cells, 150 µL of 4 µg/mL PI (Sigma-Aldrich, St. Louis, MO, USA) was added, and after brief (1–2 min) incubation at RT, 0.8 mL of water was added. All samples were kept on ice. A total of 100,000 cells were counted at each experimental point using a NovoCyte Flow cytometer (Agilent, Santa Clara, CA, USA). DHR-stained cells were detected using 488 nm for excitation and 530/30 nm for emission.

PI-stained cells were detected using 488 nm for excitation and 572/28 nm for emission. All assays were repeated three times, and the mean results are presented.

### 2.4 K^+^ efflux measurement

15 ml of cell suspension prepared as in section 2.1 were placed in a thermostated cell (30°C) with a magnetic stirrer. To measure the K^+^ concentration, an ELIS 121K K^+^-selective electrode (NV-LAB, Moscow, Russian Federation) connected to Hanna pH211 pH meter/conductometer (Hanna Instruments, Woonsocket, RI, USA) was used; the reference electrode was filled with 1 M LiNO_3_. Readings were taken every 15 minutes. To measure K^+^ concentration in the presence of glucose, 0.75 ml of 2 M glucose was added to 15 ml of cell suspension in water.

### 2.5 Glutathione monoethyl ester (GME) assay

2 mkL or 4 mkL of a 250 mM solution of GME (Sigma-Aldrich, St. Louis, MO, USA) in water was added to 50 mkL of cell suspension prepared as in section 2.1. The samples were incubated in a ThermoMixer for 1 h at 30°. 1 mkL of 0.5 mg/mL DHR in DMSO was added to the samples. The samples were incubated in a ThermoMixer for 1 h at 30°. The cells were pelleted at 13,000× g for 1 min, and the supernatant was discarded. The cells were washed once with 0.5 mL of water, and the cell pellet was suspended in 50 mkL water. Another 50 mkL of cell suspension with GME added but without the addition of DHR was subjected to all the above steps to obtain cells for propidium iodide (PI) staining. 2 mkL and 4 mkL of 250 mM GME were again added to the samples. 2.5 mkL of 2 M glucose solution was added to all tubes. The samples were incubated in a ThermoMixer at 30 °C for 1 h. Samples were then stained with PI and prepared for flow cytometry as in section 2.3.

### 2.6 Nitro-L-arginine methyl ester (L-NAME) assay

50 mkL of cell suspension from section 2.1 was pelleted and suspended in 50 mkL of 10 mM or 50 mM L-NAME solution in water (Sigma-Aldrich, St. Louis, MO, USA). The samples were incubated in a ThermoMixer for 1 h at 30°. 1 mkL of 0.5 mg/mL DHR in DMSO was added to the samples. The samples were incubated in a ThermoMixer for 1 h at 30°. The cells were pelleted at 13,000× g for 1 min, and the supernatant was discarded. The cells were washed once with 0.5 mL of water, and the cell pellet was suspended in 50 mkL water. Another 50 mkL of cell suspension after incubation with L-NAME but without the addition of DHR was subjected to all the above steps to obtain cells for propidium iodide (PI) staining. 2.5 mkL of 2 M glucose solution was added to all tubes. The samples were incubated in a ThermoMixer at 30 °C for 1 h. Samples were then stained with PI and prepared for flow cytometry as in section 2.3.

### 2.7 Statistical analysis

All measurements were carried out at least 3 times. Data are presented as mean ± SEM. One-way ANOVA statistics were calculated using the Prism 9 software (GraphPad, Boston, MA, USA).

## 3. Results and Discussion

### 3.1 ROS generation

We have previously shown that SICD occurs in cells with increased ROS content [15] and does not depend on the functional state of the mitochondrial respiratory chain [16]. The question arose about the origin of ROS in the development of SICD. To answer this question, several strains of S. cerevisiae BY4742 with deleted genes encoding ROS-generating enzymes were tested (Table 1). ΔLPD1, ΔNDE1, ΔSDH1 – with deleted mitochondrial dehydrogenases. ΔPOX1 – with deleted putative acyl-CoA oxidase involved in fatty acid beta-oxidation and localized in the peroxisomal matrix. For correct comparison, ΔAFO1 (mitochondrial ribosomal protein of the large subunit) and ΔYNO1 (member of a superfamily NOX enzymes) strains were rechecked. ΔFRE8 strain (iron/copper reductases which have similarity to YNO1) was also tested. As can be seen from Fig. 1, the deletion of these genes led to a significant drop in the number of cells with excess ROS in three strains: ΔAFO1 by 25% (p <0.05), ΔPOX1 by 29% (p <0.001) and ΔYNO1 by 35% (p <0.0001).

**Fig. 1.**
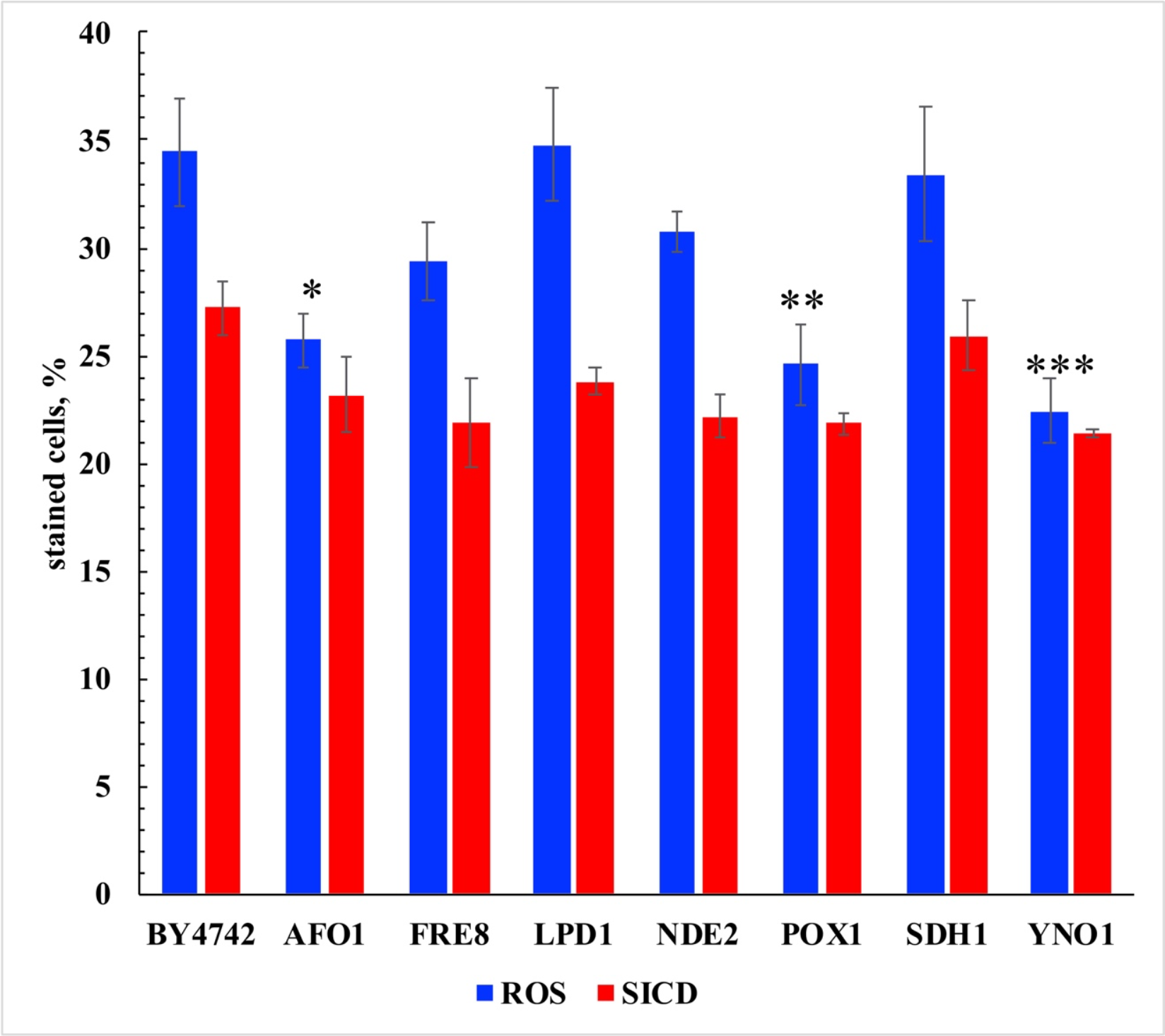
Percentage of DHR-positive cells with excess ROS and PI-positive cells with SICD in strains with impairments in ROS-generating systems after 1 hour of incubation with 100 mM glucose. Individual strains were compared to parental strain BY4742 using a one-way ANOVA test. P-value indicated as ^*^ p <0.05; ^**^ p < 0.001; ^***^ p <0.0001.

Interestingly, all three enzymes are localized in different organelles: mitochondria, vacuole, and ER, respectively. However, these differences in the number of cells with ROS did not lead to a significant change in the level of SICD - one-way ANOVA analysis showed no significant difference (p>0.05) when comparing all mutants with the parental strain BY4742. This observation contradicts the fact that in yeast at the stationary stage of growth, mutants with a defective mitochondrial respiratory chain show greater sensitivity to SICD [9]. In addition, the generation of ROS turned out to be not as tightly linked to the development of SICD as previously assumed and allows fluctuations within certain limits. It can be assumed that the generation of ROS by Afo1p, Pox1p, and Yno1p is not the main mechanism for the production of free radicals involved in SICD. Therefore, deletion of these genes caused a decrease in the number of cells with excess ROS but did not significantly reduce the number of dead cells.

To distinguish these fluctuations from significant changes in ROS content and SICD values, experiments were carried out on the effect of exogenous antioxidants. Ascorbate and glutathione suppress the generation of ROS and the development of SICD [15]. However, both of these compounds must be in neutralized form at a pH close to 7 to effectively enter the cell. However, the neutral pH of the incubation medium itself very effectively suppresses both processes under consideration [16]. To evaluate the actual effect of antioxidants, the SICD assay was carried out in the presence of GME. GME is a cell permeable compound and can therefore be used in an unbuffered environment. GME at concentrations of 10 and 20 mM effectively suppressed both ROS generation (by 44% at 20 mM) and SICD (by 46% at 20 mM) (Fig. 2a). This fact suggests that ROS generation and SICD development are functionally related.

**Fig. 2.**
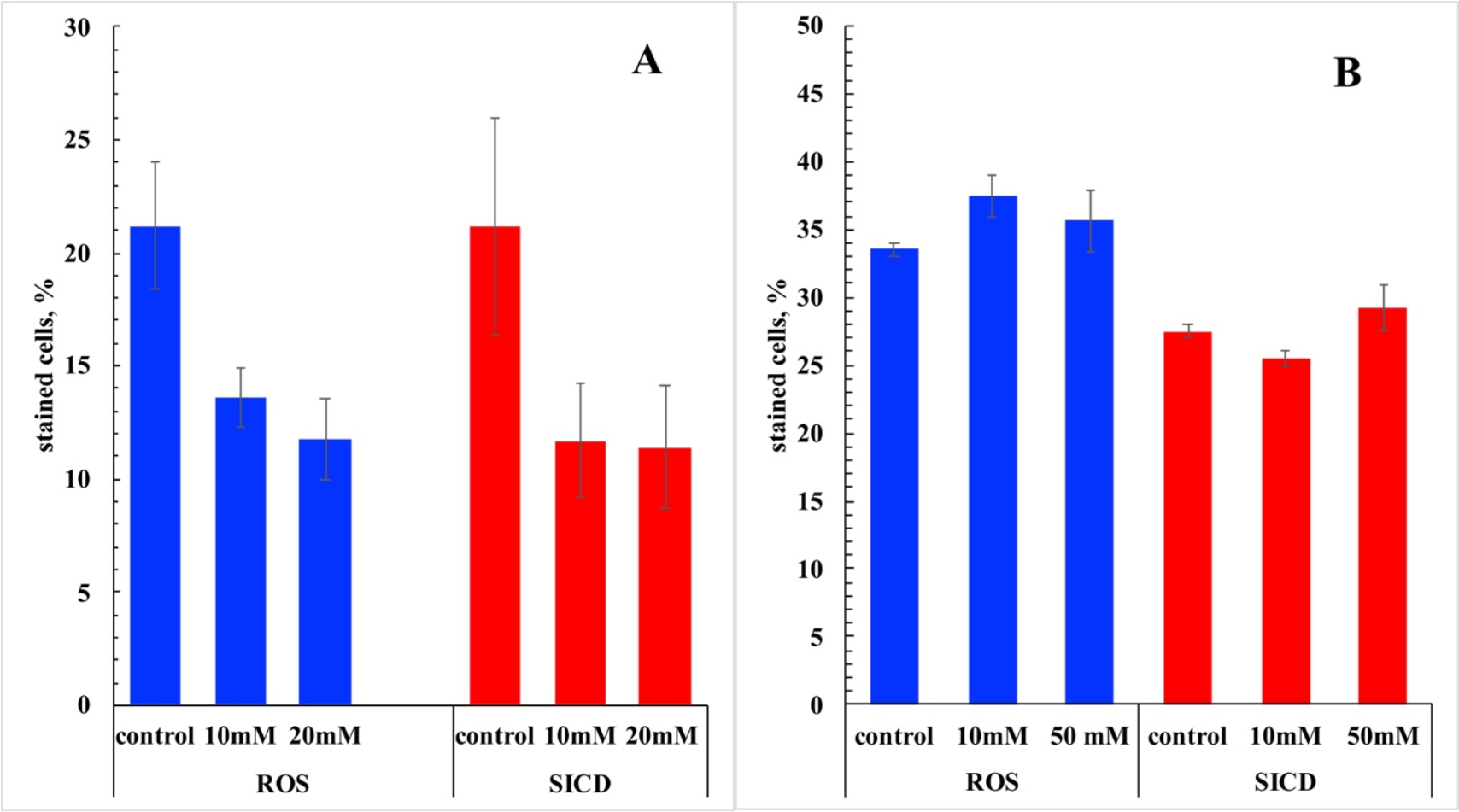
Effect of (**A**) GME (ROS scavenger) and (**B**) L-NAME (NRS scavenger) on the percentage of DHR- and PI-positive cells in strain BY4742 after 1 hour of incubation with 100 mM glucose. Data are presented as mean±SEM from 3 independent experiments.

There is another important aspect to consider when analyzing ROS generation. The DHR fluorescent probe used to detect ROS is not specific, and may also react to NRS (Nitrogen Reactive Species) [18]. To test this assumption, a SICD assay of BY4742 cells was carried out in the presence of L-NAME (NG-nitro-L-arginine methyl ester is a nonspecific inhibitor of nitric oxide synthase). As can be seen from Fig. 2b, L-NAME did not affect ROS generation and SICD development at concentrations of 10 mM and 50 mM. At higher concentrations, L-NAME showed even a weak pro-oxidant effect (not illustrated). Thus, it can be concluded that NRS is not involved in SICD and that it is ROS that oxidizes DHR to its fluorescent derivative, rhodamine.

Strains ΔTSA1 (thioredoxin peroxidase which acts as both ribosome-associated and free cytoplasmic antioxidant) and ΔYDR034W-B (protein whose biological role is unknown and contains conserved transmembrane CYSTM module) were used to test the effect of endogenous antioxidants on ROS generation and SICD development. It was expected that deletion of these genes would lead to an increase in the number of cells with excess ROS and, therefore, to an increase in the number of dead cells. But knock-out mutants for these genes showed ROS and SICD values close to BY4742 – p <0.05 (not illustrated). It is interesting to note that while TSA1 is involved in the cellular response to oxidative stress at the cytoplasmic level, the CYSTM motif YDR034W-B is thought to neutralize ROS in the lipid bilayer [19]. The results are paradoxical - ROS that cause SICD are inaccessible to TSA1 in the cytosol and are not localized in the membrane. One explanation is that TSA1 has a paralogue, TSA2, that successfully neutralizes some of the ROS in the ΔTSA1 strain.

The above data provide evidence that the well-known major intracellular producers of ROS are not involved in the development of SICD in exponentially growing cells. NRS also does not participate in SICD. At the same time, the almost 2-fold suppression of SICD by GME indicates that ROS is a trigger for SICD. This fact also excludes the possibility that the observed increase in the number of cells with ROS is a result of oxidative stress and is not related to SICD. In this case, it would be possible to decouple the ROS generation from the SICD. Another evidence in favor of the conjugation of the processes of ROS generation and the development of SICD is the fact that we previously discovered the simultaneous suppression of these phenomena by extracellular neutral pH by more than 90% [16]. When incubated with glucose in 50 mM HEPES buffer pH 7.0, all of the above strains showed an almost complete absence of ROS and SICD. 150 mM extracellular KCl also suppressed ROS generation and SICD development by more than 85% [16]. It can be concluded that ROS leading to SICD is generated by some unknown enzymes/processes that are sensitive to ΔpH and ΔΨ. Another group of researchers came to the same conclusion when studying SICD in yeast at the stationary growth stage [20].

### 3.2 Transport systems

Transport of nutrients across the plasma membrane and organelle membranes depends on membrane potential and/or ΔpH. Both of these parameters regulate the generation of ROS and SICD under our conditions [16]. This indicates the possible (indirect, through a change in ΔΨ) involvement of transport systems in the phenomenon under study. Strains with knockout genes TRK1, TRK2, AGP1, GAP1, and PHO84, the main transport systems for the uptake of K^+^, amino acids and inorganic phosphorus from the environment, were tested (Fig.3). ROS generation and SICD levels in strains ΔAGP1, ΔGAP1, and ΔPHO84 were not significantly different compared to the parental strain BY4742 (p > 0.05), therefore these transport systems are not involved in SICD.

**Fig. 3.**
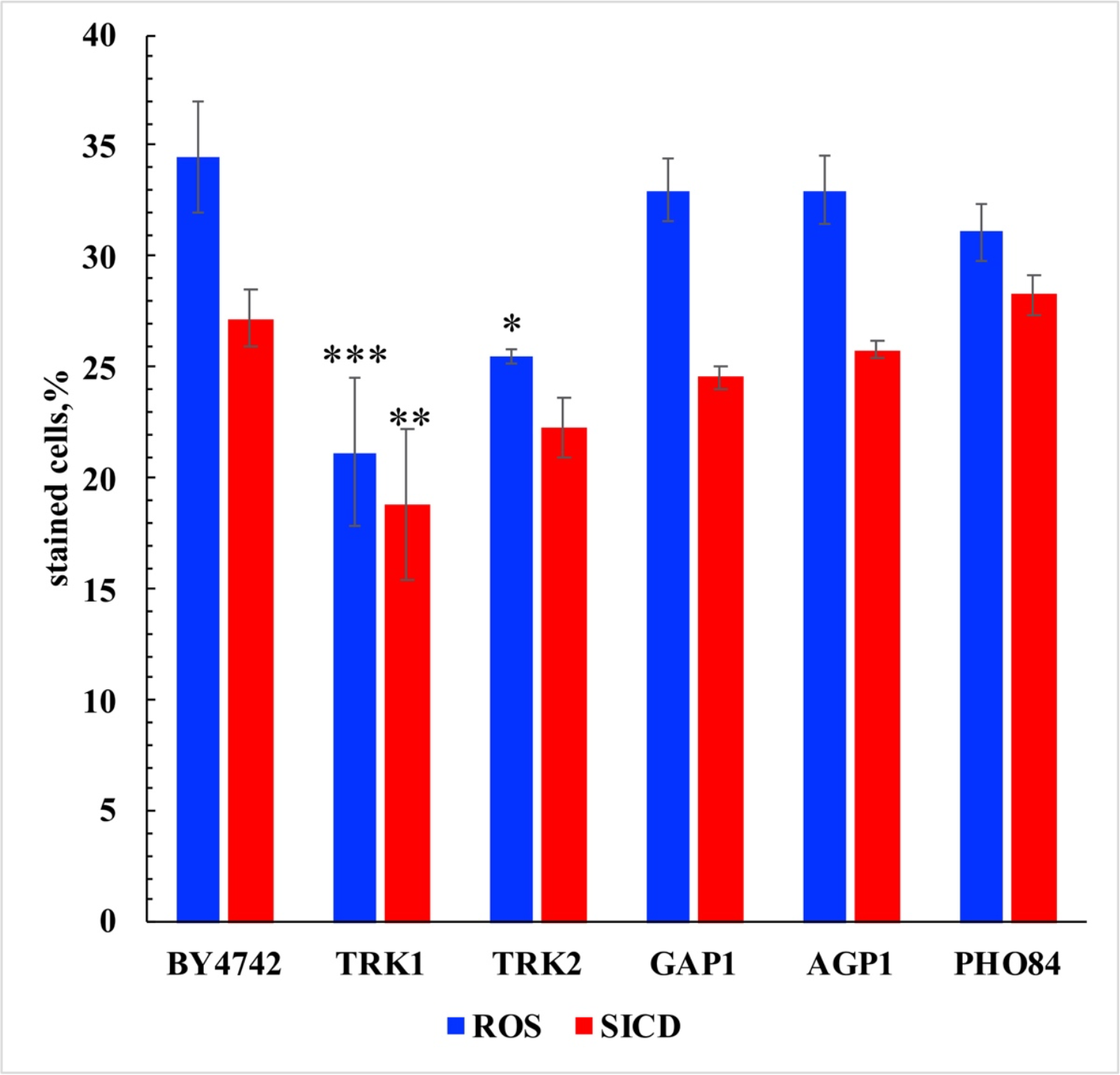
Percentage of DHR- and PI-positive cells in strains lacking transport systems after 1 hour of incubation with 100 mM glucose. Individual strains were compared to parental strain BY4742 using a one-way ANOVA test. P-value indicated as ^*^ p <0.05; ^**^ p < 0.001; ^***^ p <0.0001.

ΔTRK1 and ΔTRK2 showed a significant decrease in the number of cells with excess ROS compared to BY4742. ΔTRK1 by 39% (p <0.0001) and ΔTRK2 by 26% (p <0.05). However, only ΔTRK1 showed a significant mitigation in SICD development by 31% (p < 0.001). The findings show a fundamental difference in SICD between exponentially growing and stationary yeast. TRK2 plays a critical role in the survival of SICD in stationary growth phase cells [21], whereas ΔTRK1 was more resistant under our conditions. In ΔTRK1 and ΔTRK2, ROS generation and SICD showed lower sensitivity to 150 mM KCl than BY4742 (Table 2).

**Table 2.** Effect of 150 mM KCl on the percentage of cells with excess ROS and the percentage of dead cells.

| strain | ROS |  | SICD |  |
| --- | --- | --- | --- | --- |
|  | glucose | glucose + KCl | glucose | glucose + KCl |
| BY4742 | 34,5 | 4,5 (87%) | 27,2 | 5,9 (78%) |
| ΔTRK1 | 21,1 | 5,6 (73%) | 19 | 7,1 (62%) |
| ΔTRK2 | 26 | 6,3 (76%) | 22 | 6,4 (71%) |

The data shows the percentage of DHR-positive and PI-positive cells after 1 hour of incubation with 100 mM glucose ± 150 mM KCl. The percentage of inhibition by 150 mM KCl is indicated in parentheses.

ΔTRK1 lacks a major K^+^ transporter, whereas ΔTRK2 lacks a minor transporter. This explains the difference in the suppression of the generation of ROS and SICD by extracellular K^+^ in these two strains. That is, to suppress the generation of ROS and the development of SICD, it is important not only the dissipation of ΔΨ by exogenous K^+^ but also the entry of K^+^ into the cell. The role of K^+^ in the development of SICD is not yet clear enough. For example, BY4742 cells release almost half as much intracellular K^+^ when incubated with glucose as when incubated in water (Fig. 4).

**Fig. 4.**
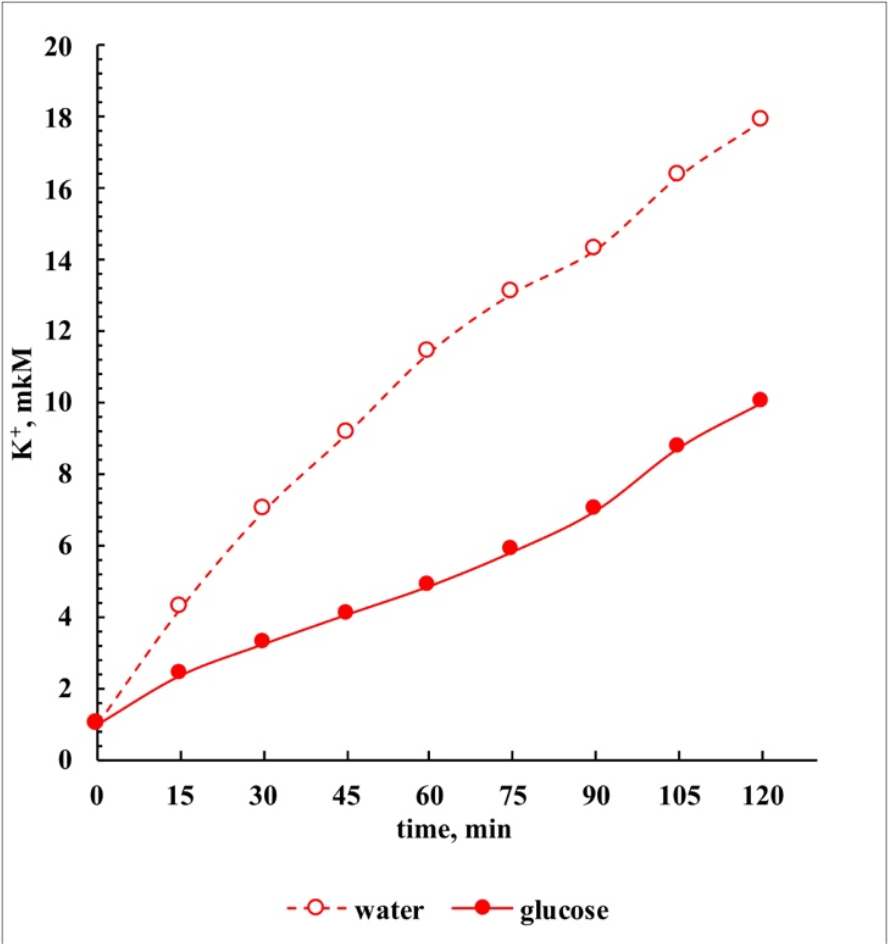
Dynamics of the efflux of intracellular K^+^ into the medium during incubation of yeast BY4742 in water or in the presence of 100 mM glucose.

Apparently, the reverse absorption of K^+^ occurs due to the work of the K^+^/H^+^ antiporter. This is confirmed by the fact of stronger acidification of the medium during incubation with glucose in the presence of 150 mM KCl [16]. This leads to the fact that the content of intracellular K^+^ should not drop significantly. This assumption is in good agreement with the hypothesis that SICD is a consequence of vacuole collapse after loss of K^+^ [20]. Potassium entering through the K^+^/H^+^ antiport apparently accumulates again in the vacuole, thereby preventing its collapse. But then it is not clear why BY4742 cells, losing twice as much K^+^ when incubated in water, remain alive. Induction of SICD by loss of K^+^ from the vacuole, however, does not explain the suppression of SICD by neutral pH. Neutral pH and exogenous K^+^ lead to the dissipation of ΔΨ [16]. Therefore, the ΔΨ status is decisive in the initiation of SICD and ROS generation.

### 3.3 Genes expressed in the S-phase of the cell cycle

SICD in exponentially growing yeast affects only cells in the S-phase of the cell cycle [15]. Table 1 presents some of the genes expressed in the S-phase that have been tested for their ability to modify SICD. To select genes, we used the Yeast Protein Database, which is currently unavailable. An alternative database can be CycleBase (https://cyclebase.org/CyclebaseSearch accessed 15 September 2023). Analysis of the database allowed the selection of several genes that might be involved in SICD. We selected the following genes. YNR066C, PRM5, and YET1 - proteins whose biological role is unknown. SEO1 and WSC2 - transmembrane proteins. PRB1 - serine-type endopeptidase is involved in vacuolar protein catabolism. ATG41 – a protein involved in mitophagy, and macroautophagy. STR3, MMP1, SAM3, MET1, MET5, MET10, MET14 and MET28 – proteins involved in sulfur amino acid metabolism. Interestingly, many genes involved in sulfur metabolism are activated in the S-phase of the cell cycle. We included these genes in the list because SICD under our conditions was accompanied by accumulation in the cell and release of octanoic acid into the medium [17]. Octanoic acid is a toxic product. After the addition of two sulfur atoms, it is converted into lipoic acid, an important cofactor in many biochemical reactions. We hypothesized that under conditions leading to SICD, some stages of sulfur assimilation are disrupted. The results obtained are presented in Fig.5.

**Fig. 5.**
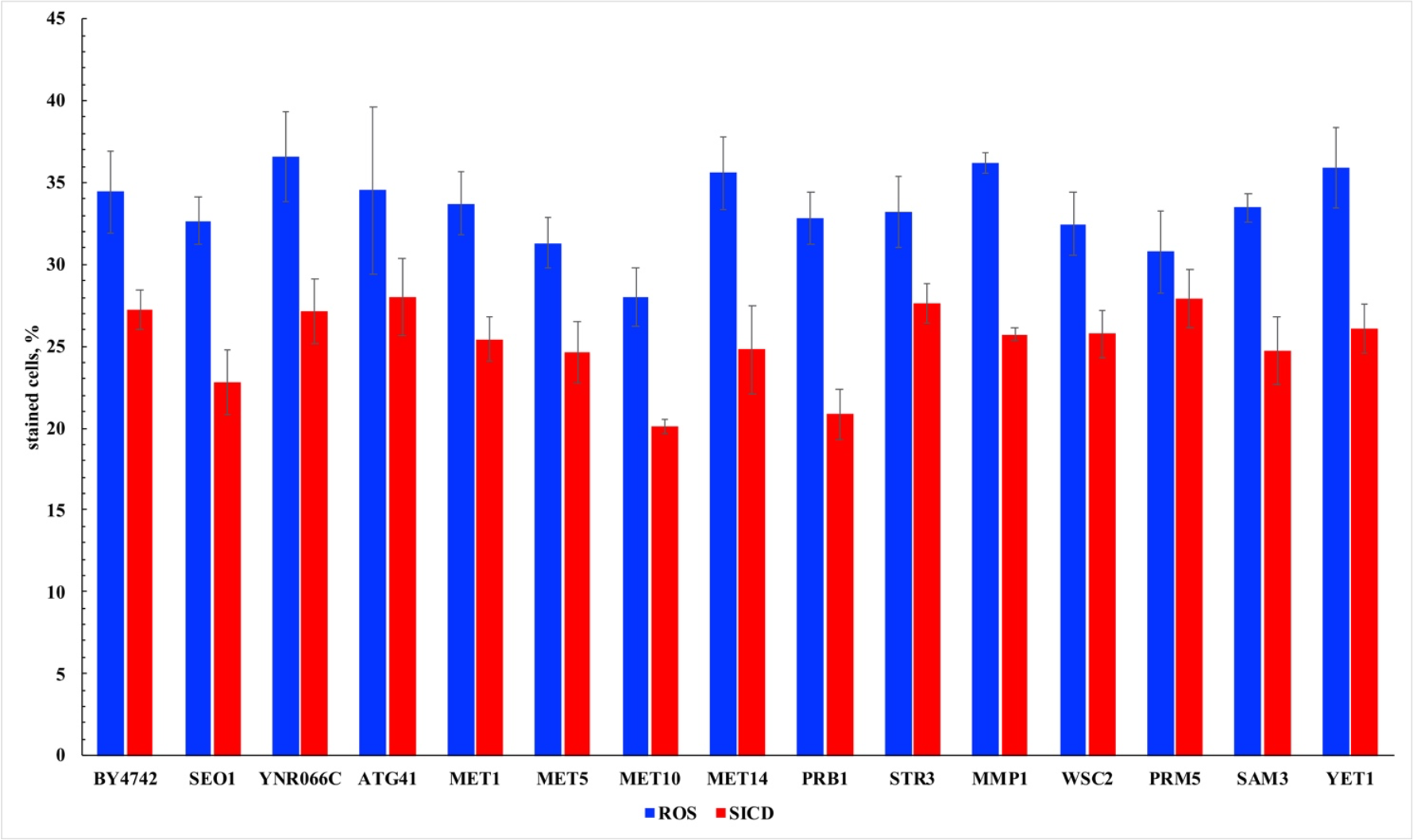
Percentage of DHR-positive and PI-positive cells in strains with deleted genes expressed in the S-phase of the cell cycle, after incubation with 100 mM glucose for 1 hour. One-way ANOVA test showed a p-value > 0.05 for ROS generation and SICD development in all strains when compared individually with the parental strain BY4742.

None of the deletion strains tested showed a significant difference in ROS generation and SICD development compared to parental BY4742 (p > 0.05). It should be noted that such a distinctive characteristic as the mitigation of ROS generation and SICD development by neutral pH or 150 mM KCl was completely reproduced in all of the above strains. It can be concluded that the tested genes expressed in S-phase are not involved in either the generation of ROS or the development of SICD.

### 3.4 Effect of yeast homologs of SMT15

In 2014, a group of researchers discovered an interesting fact that allows us to establish a new functional relationship between the phases of the cell cycle and the amount of glutathione in Chlamydomonas reinhardtii [22]. Deletion of the detected gene SMT15 (encoding putative membrane-localized sulfate/anion transporters) resulted in the fact that the level of glutathione in the cell did not fall in the S- and M-phases of the cell cycle. Moreover, ΔSMT15 had an increased reduced-to-oxidized glutathione redox ratio throughout the cell cycle. In the yeast S. cerevisiae, the homologous gene is VSB1, a predicted integral membrane protein whose biological role is unknown. But recently, the product of this gene was characterized as a vacuolar membrane protein required for the uptake and storage of arginine in nitrogen-replete cells [23]. In addition, a loose search for SMT15 homologs in BLASTN finds another yeast homolog, YPR003C (distantly related to putative sulfate permease) [24]. The results are presented in Fig.6.

**Fig. 6.**
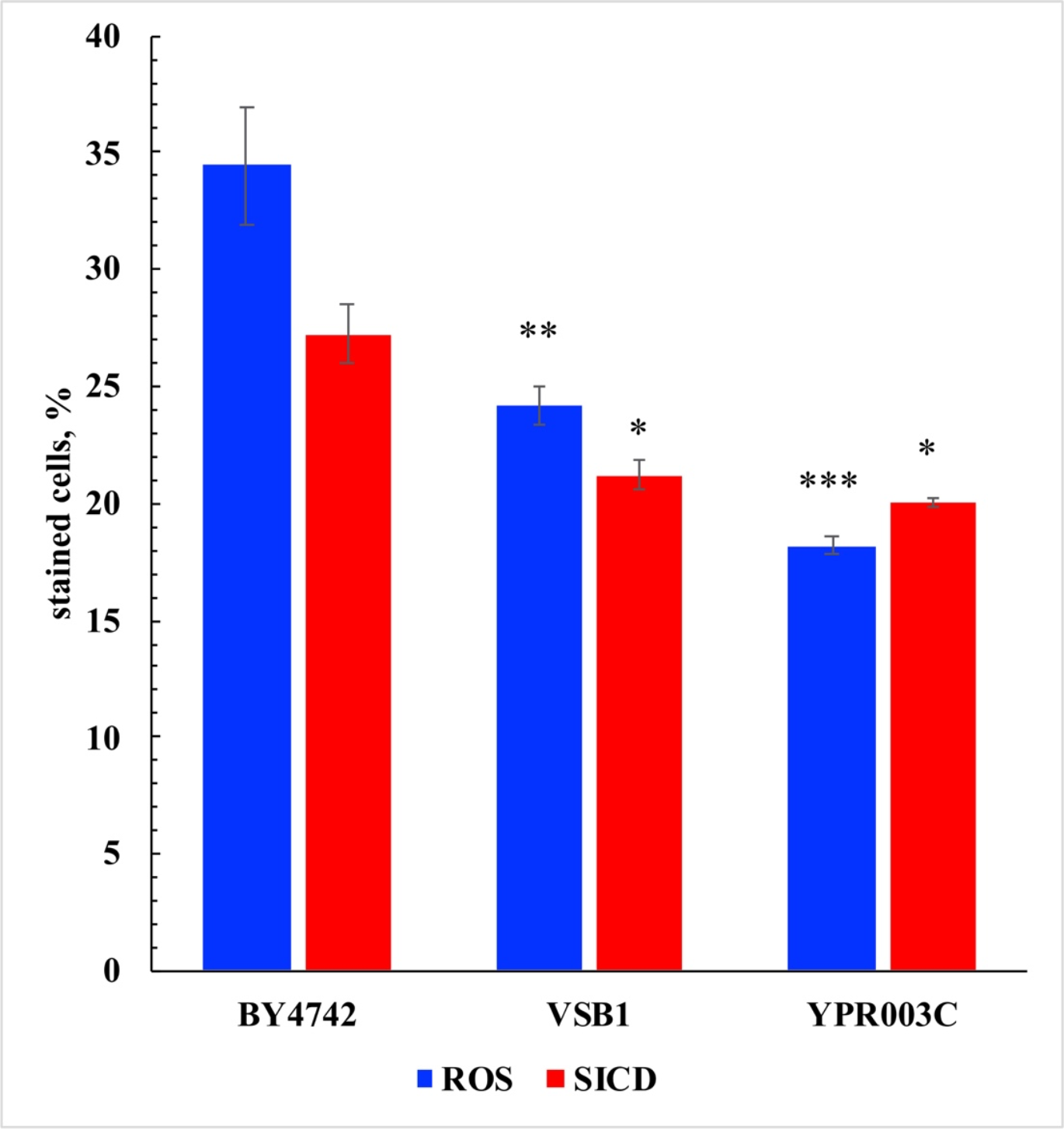
Percentage of DHR-positive and PI-positive cells in strains ΔVSB1 and ΔYPR003C after incubation with 100 mM glucose for 1 hour. The strains were compared with the parental strain BY4742 using a one-way ANOVA test. P-value indicated as ^*^ p <0.05; ^**^ p < 0.001; ^***^ p <0.0001.

Both strains showed a significant decrease in the number of cells with excess ROS – p <0.001 for ΔVSB1 and p <0.0001 for ΔYPR003C. YPR003C is a sulfate permease and therefore its deletion had a more pronounced effect, which is consistent with the assumption of the role of glutathione, the synthesis of which requires sulfur. YPR003C is predicted to localize to the endoplasmic reticulum (https://www.yeastgenome.org/locus/S000006207) and may be involved in creating sulfate deficiency in the cytoplasm. I.e., if deletion of these two genes leads to an increase in intracellular glutathione levels (similar to ΔSMT15 in Chlamydomonas reinhardtii), then the decrease in the number of cells with excess ROS has a very convincing explanation. The number of cells with ROS decreased in ΔVSB1 by 30%, and in ΔYPR003C by 47%. Both strains also showed a decrease in the number of cells with SICD, although not as significant - p < 0.05 for both strains (22% in ΔVSB1, and 27% in ΔYPR003C). Data obtained from these strains again show a loose relationship between the number of cells with excess ROS and the number of dead cells. At the same time, these data confirm the role of glutathione, which maintains the reducing potential in the cell, in the suppression of SICD. There is an interesting observation that can be applied to the analysis of ROS generation, SICD, and cell cycle stages. It has been shown that the cellular redox potential undergoes cyclic changes [25]. Moreover, the S- and M-phases of the cell cycle occur in the reductive phase, and cell division itself occurs in the oxidative phase. However, most of the cellular machinery involved in DNA synthesis is expressed in the oxidative phase. An important note must be made here - the data were obtained on a chemostat culture under conditions of partial synchronization of the culture both in the phases of the cell cycle and in the yeast metabolic cycle (YMC) [25]. Due to the fundamental difference between batch and continuous culture, it can be assumed that under our conditions, a YMC shift may occur, resulting in the S-phase falling at the oxidative stage of YMC. As a result, ROS generated (via an unknown metabolic pathway) cannot be completely neutralized by glutathione, the peak production of which occurs during the reductive phase of YMC [26].

## 5. Conclusions

It was shown that in strains ΔAFO1, ΔPOX1, and ΔYNO1, in which the mechanisms of ROS generation in organelles such as mitochondria, peroxisome, and ER are impaired, there is a significant decrease in the number of cells with excess ROS (by 25%, 29% and 35%, respectively). However, this decrease is not accompanied by a significant decrease in the number of cells with SICD. GME, unlike L-NAME, showed a significant mitigation of ROS and SICD generation. However, the endogenous antioxidants TSA1 and YDR034W-B did not have a significant effect on the studied processes. A decrease in the number of cells with ROS and SICD in the ΔTRK1 and ΔTRK2 strains showed a partial dependence of SICD on ΔΨ but did not confirm the key role of ΔTRK2 in the suppression of SICD, as was shown in stationary cells [21]. The PM voltage-dependent transport systems AGP1, GAP1, and PHO84 were not involved in the regulation of SICD. Thus, the critical dependence of SICD on ΔΨ remains unexplained. . It was also shown for the first time that genes expressed in the S-phase of the cell cycle (in which SICD develops) are not involved in the development of SICD. Strains ΔVSB1 and ΔYPR003C, with disturbances (presumably) in sulfur metabolism, showed a significant decrease in the number of cells with excess ROS and with SICD. The results obtained confirm an interdependence, although not a strict one, between ROS accumulation and the development of SICD. In all tested strains, neutral pH and 150 mM KCl decreased the number of cells with excess ROS and with SICD to the same extent as in the parent strain BY4742 – almost completely.

